# An Event-related Potential Comparison of Facial Expression Processing between Cartoon and Real Faces

**DOI:** 10.1101/333898

**Authors:** Jiayin Zhao, Yifang Wang, Licong An

## Abstract

Faces play important roles in the social lives of humans. In addition to real faces, people also encounter numerous cartoon faces in daily life. These cartoon faces convey basic emotional states through facial expressions. Using a behavioral research methodology and event-related potentials (ERPs), we conducted a facial expression recognition experiment with 17 university students to compare the processing of cartoon faces with that of real faces. This study used face type (real vs. cartoon) and participant gender (male vs. female) as independent variables. Reaction time, recognition accuracy, and the amplitudes and latencies of emotion processing-related ERP components such as N170, vertex positive potential (VPP), and late positive potential (LPP) were used as dependent variables. The ERP results revealed that cartoon faces caused larger N170 and VPP amplitudes as well as a briefer N170 latency than did real faces; that real faces induced larger LPP amplitudes than did cartoon faces; and that angry faces induced larger LPP amplitudes than did happy faces. In addition, the results showed a significant difference in the brain regions associated with face processing as reflected in a right hemispheric advantage. The behavioral results showed that the reaction times for happy faces were shorter than those for angry faces; that females showed a higher facial expression recognition accuracy than did males; and that males showed a higher recognition accuracy for angry faces than happy faces. These results demonstrate differences in facial expression recognition and neurological processing between cartoon faces and real faces among adults. Cartoon faces showed a higher processing intensity and speed than real faces during the early processing stage. However, more attentional resources were allocated for real faces during the late processing stage.

---

Faces play important roles in human social life. They convey unique identity information and basic emotions through facial expressions. Facial expression refers to the various emotional states that people present through both the automatic and intentional control of the eye, facial, and oral muscles. In daily life, facial expressions provide important non-verbal forms of information and communication (Batty & Taylor, 2003). As a non-verbal signal, facial expressions are an important basis and prerequisite for emotional communication as well as the generation of emotional experience. The ability to recognize a facial expression reflects an individual’s ability to infer the psychological states of others through emotional expressions (Nelson, 1979). Facial expression recognition not only helps to determine internal emotional states (Thompson & Meltzer, 1964) and the intentions conveyed by an individual but also provides feedback and induces social interactions (Erickson & Schulkin, 2003). Ekman and Friesen (1978) summarized six basic human facial expressions including happiness, sadness, surprise, fear, anger, and disgust. These facial expressions have been identified and confirmed across different cultural contexts (Ekmm & Friesen, 1971).

With the development of social communication, cartoons have entered people’s lives. In addition to real faces, people also encounter many cartoon faces on daily basis. Moreover, common social networks (e.g., WeChat) provide various cartoon face emoji for communicating and expressing emotions. Compared with real faces, cartoon faces usually have larger eyes, smaller noses, and finer skin texture (Schindler, Zell, Botsch & Kissler, 2017). Chen and colleagues (2010) found that people developed a preference for real faces with larger eyes after adaption to cartoon faces with unusually large eyes in Japanese cartoons. Some researchers compared cartoon faces and real faces with regard to recognition accuracy and reaction time. Kendall, Raffaelli, Kingstone, and Todd (2016) asked participants to identify emotions on five sets of briefly presented faces that ranged from photorealistic to fully iconic. The results showed stronger emotion recognition accuracy for cartoonized faces. In another study (Wang, Wang, Wang & Lu, 2012), participants showed faster reaction times to real faces than cartoon faces when they were required to determine whether an image was a face or a car. However, research on the recognition of cartoon and real faces has shown mixed results. Using synthesized emotion images, Hoptman and Levy (1988) studied the processing preference of left- and right-handed individuals for cartoon and real faces. The results failed to reveal significant difference between cartoon and real faces.

Both cartoon and real faces convey emotional information through facial expression. The six basic facial expressions can be categorized as positive or negative expressions. Mixed results have been reported by research on the reaction times and recognition accuracies of positive and negative facial expressions. Some believe that reaction times for positive expressions are faster than those for other facial expressions. Eimer, Holmes and Mcglone (2003) found that recognition of happiness is faster than that of other basic facial expressions. In an identification task regarding whether facial expressions were neutral or emotional, participants showed the shortest reaction time for happiness, the longest reaction times for sadness, the lowest error rate for surprise, and the highest error rate for sadness. Calvo and Lundqvist (2008) asked participants to press different buttons for each of the six basic facial expressions and found similar results. However, they found that the recognition accuracy for happiness was highest, whereas the recognition accuracy for fear was lowest. Other studies have suggested people recognize negative facial expressions faster than positive ones. Hansen and Hansen (1988) found that the search speed for angry face targets was faster when both angry and happy faces served as targets and distractors. Eastwood, Smilek and Merikle (2001) reported that the search for negative facial expressions (sadness) was faster than positive facial expressions (happiness) when neutral faces were used as distractors.

The above experiments investigated the differences among different facial types and facial expressions from the perspective of behavioral science. Using this behavioral research as a basis, other researchers have also used event-related potentials (ERPs) to study the neurophysiological basis behind these differences. ERPs refer to the changes in the electrical potential of various brain regions when a stimulus is applied or removed to the sensory system or a certain part of the brain (Wei & Luo, 2002). ERPs directly reflect electrical neurological activity. ERPs have been widely applied in face processing research because it provides high temporal resolution, real-time and non-invasive measurement, and a connection among stimulus events, psychological reactions, and brain activity. ERPs can be used to classify different visual stimulus and differentiate disparate emotional states. Without any participant response, ERP testing enables the measurement of emotional attitudes that people are unwilling to express (Bernat, Bunce & Shevrin, 2001).

The ERP components related to faces and facial expressions include N170, vertex positive potential (VPP), late positive potential (LPP), and others. N170 is primarily distributed in the occipito-temporal region of the brain and usually shows a larger response in the right hemisphere (Rossion & Jacques, 2008). N170 is a face-specific ERP component, and its peak shows face selectivity. N170 is only induced by face stimuli (i.e., not by furniture, cars, hand gestures, or other stimuli)(Bentin, Allison, Puce & McCarthy, 1996). Related to face type, research has shown that the N170 components induced by real and cartoon faces do not significantly differ (Wang et al., 2012). Another study (Sagiv & Bentin, 2001) showed that real faces induced a stronger N170 effect than abstract sketches of faces. Compared with schematic faces, however, the difference was not significant. Facial expressions are also related to N170 during early processing (Galli, Feurra & Viggiano, 2006). A meta-analysis revealed that larger N170 amplitudes are associated with facial expressions of anger, fear, and happiness compared with neutral facial expressions (Hinojosa, Mercado & Carretié, 2015). Rellecke, Sommer, and Schacht (2012) required participants to explicitly or implicitly process happiness, anger, and neutral faces. Their results showed that emotional faces induced larger N170 amplitudes than did neutral faces under both processing conditions. With respect to different facial expressions, Batty and Taylor (2003) recorded the ERPs of participants responding to the six basic facial expressions and neutral expressions. The results showed that positive expressions resulted in shorter N170 latencies than negative expressions and that fear expressions induced significantly larger amplitudes than did other expressions.

N170 has a corresponding positive component at the mind-central sites, namely VPP. VPP and N170 have similar functional properties. They are two manifestations of the same brain processes (Joycea & Rossion, 2005). VPP sometimes shows more sensitivity to facial expression information than N170, and VPP is influenced by facial expressions when N 170 is not (Ashley, Vuilleumier & Swick, 2004).

Additional processing of emotional expression is reflected by the LPP component (Bublatzky, Gerdes, White, Riemer & Alpers, 2014). LPP waves originate from the occipital lobe and the posterior parietal cortex (Keil et al., 2002), reflecting the cerebral cortex’s evaluation of emotional stimuli, working memory characterization, decision making, and response-related processing (Schupp, Flaisch, Stockburger & Junghofer, 2006). LPP waves are sensitive to various emotional stimuli including faces (Flaisch, Häcker, Renner & Schupp, 2011; Schindler & Kissler, 2016; Schupp, Junghöfer, Weike & Hamm, 2004; Steppacher, Schindler & Kissler, 2015; Wieser, Pauli, Reicherts & Mühlberger, 2010). Using the International Affective Picture System (IAPS) to examine explicit emotional processing, researchers found that images of emotional scenes induced larger LPP amplitudes than did neutral scenes (Hajcak, Moser & Simons, 2006). However, the findings related to the influence of facial expression on LPP are not consistent. Although some reports have concluded that negative expressions (e.g., sadness) induce smaller amplitudes than do positive expressions (e.g., happiness) (Hietanen & Astikainen, 2013), others have found that negative expressions induce larger LPP components than do positive expressions (Zhu & Liu, 2014). In addition, other studies have not found significant differences between the processing of positive and negative expressions (Codispoti, Ferrari & Bradley, 2006; Recio, Sommer & Schacht, 2011). Krolak-Salmon, Fischer, Vighetto and Mauguière (2001) asked participants to view images of different facial expressions (e.g., fear, happiness, disgust, surprise, and neutral expressions) and recorded their ERPs during two different tasks with the same stimuli. First, participants were instructed to focus on the gender of the faces by counting the number of males or females. Second, they were asked to focus on facial expressions by counting the number of faces that looked surprised. The results showed significant differences between late-latency ERPs to emotional faces and those to neutral faces (between 250 and 550 ms of latency). The activation was symmetric in the occipital lobe. The ERP components of different facial expressions differed between 550 and 750 ms on the right side of the occipital cerebral region. Topographic maps of these differences showed specific right temporal activity related to each emotional expression.Differences were also found with regard to the processing of different face types. Researchers studied adults’ ERP processing of real and cartoon faces with neutral expressions (Wang et al., 2012). The results indicated that real faces induce significantly higher average LPP amplitudes in the occipital and temporal regions than do cartoon faces. The processing of the cartoon faces showed obvious lateralization, primarily in the right parietooccipital area, whereas the processing of the real faces was primarily in the parietooccipital areas of both sides. Schindler et al. (2017) employed six face-stylization levels varying from abstract to realistic and investigated the difference in the processing of real and cartoon faces. The results showed that the LPP amplitude increased as the faces became more realistic. The above studies suggest that different face types and facial expressions induce significantly different ERP components and amplitudes in different brain regions.

In addition, other studies have investigated the influence of participant gender on the recognition of facial expressions. Hoffmann and colleagues (Hoffmann, Kessler, Eppel, Rukavina & Traue, 2010)asked male and female participants to identify six basic but subtle facial emotions (50% emotional content). The results showed that women were more accurate than men at recognizing subtle facial displays of anger, disgust, and fear, suggesting that women are more sensitive to negative emotions. This result might be related to the role of women throughout evolution. The treatment of emotions comes from a corresponding neural basis. Even if the influence of gender is not reflected in the accuracy or speed of facial expression recognition, differences remain between men and women with regard to neural activity. Wildgruber, Pihan, Ackermann, Erb & Grodd (2002) found no behavioral difference between males and females with regard to differentiating happy from sad sounds. However, higher response amplitudes within the left-hemisphere posterior middle temporal gyrus were found among women compared with men, whereas a larger increase of activation within the right middle frontal gyrus was observed among the latter. Han, Gao, Humphreys & Ge (2008) found significant differences in the behaviors and brain activities between men and women during emotion-related tasks. Women showed faster threat detection times, while men showed stronger posterior parietal activation.

In summary, the existing research suggests that the N170, VPP, and LPP ERP components are closely related to facial expression processing and that each component presents different properties when processing the unique facial expressions of real faces. However, consistent conclusions do not exist regarding the comparison of processing methods, speeds, and intensities between cartoon faces and real faces. With respect to facial expression selection, happiness is usually used as a positive expression, whereas anger and sadness are usually selected as negative expressions (Schindler et al., 2017; Hansen & Hansen, 1988; Eastwood et al., 2001; Bentin et al., 1996; Hietanen & Astikainen, 2013). The present study used anger and happiness for comparison because the recognition accuracy of anger is higher than that of sadness (Eimer et al., 2003). Moreover, significant differences exist between males and females with regard to anger recognition accuracy (Calvo & Lundqvist, 2008; Hoffmann et al., 2010). The present study used an ERP methodology to investigate the processing of real and cartoon facial expressions among men and women. We hypothesized that (1) recognition time would be faster with regarding to a positive emotion (i.e., happiness) than a negative emotion (i.e., anger); (2) women would recognize facial expressions faster and more accurately than would men; (3) the late component LPP, but not N170 or VPP, would be affected by emotional valance; and (4) face type (i.e., real and cartoon faces) would influence the amplitudes and latencies of N170 and VPP as well as the amplitude of LPP.

## Method

### Participants

We recruited 17 participants (11 males, 6 females; average age = 24.18, SD = 2.32) from universities in Beijing. All participants were right-handed, had normal hearing and vision (with or without correction), and no history of hearing, neurological, or psychiatric disorders. Participants signed an informed consent document prior to the experiment and were compensated for their time after the experiment. This study was approved by the ethical committee of Beijing Key Laboratory of Learning and Cognition of Capital Normal University.

### Materials

The pictures used in the experiment were selected from the Chinese Facial Affective Picture System (CFAPS; Wang and Luo, 2005) and the Japanese Female Facial Expression (JAFEE) database. Fifty pictures of happy faces (25 males and 25 females) and 50 pictures of angry faces (25 males and 25 females) were selected from the two picture databases. In total, 100 pictures were selected. We used MYOTee (a cartoon image editor) to convert these faces into cartoon faces. Subsequently, we used Photoshop to overlay the cartoon faces onto the original pictures for fine-tuning, and we retained the same face structure and hairstyle to synthesize 100 cartoon facial expression pictures. In total, 200 pictures were used in this experiment. All pictures were presented in black and white with a resolution of 260 × 300 at a consistent contrast.

**Table 1.**
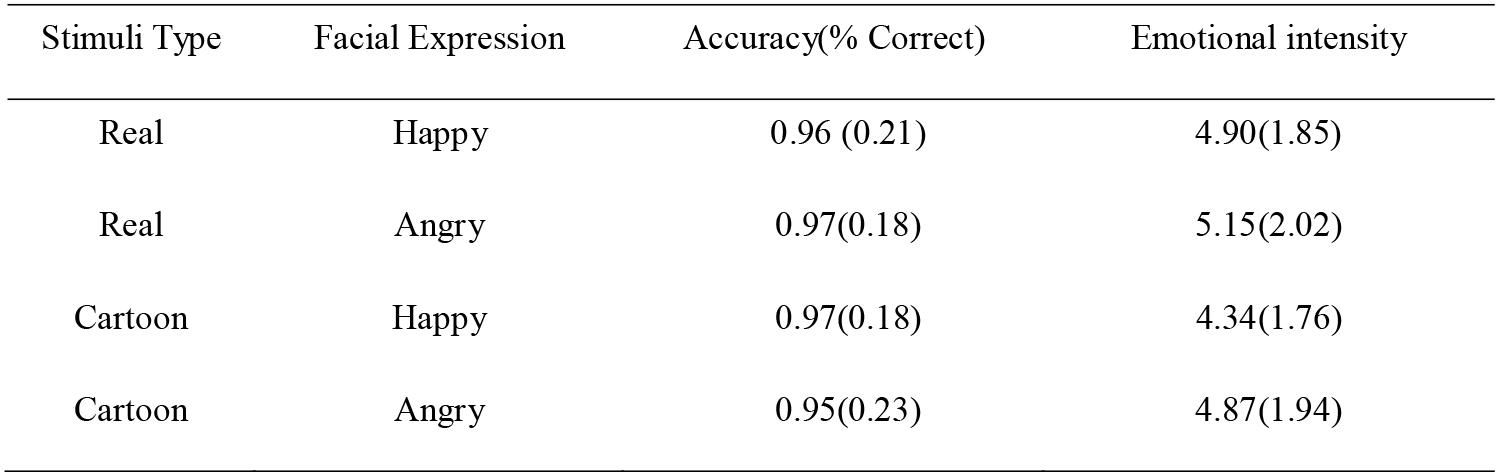
Means (and Standard Deviations) for Accuracy of Facial Expression Recognition and Valuation of Emotional Intensity

Twenty additional volunteers (non-participants; mean age = 25.3 years) evaluated the pictures. The evaluation included the identification of facial expression type (i.e., by pressing the “G” key for happiness and the “F” key for anger) and a Likert rating of the facial emotion (9 = extremely happy or angry; 1 = not at all happy or angry). The evaluation results revealed a recognition accuracy of 95.9%, with an emotion intensity rating of 4.81 ± 1.91 (Table 1). Therefore, all 200 pictures were retained as stimuli for the experiment.

### Procedure

The experiment was conducted in a quiet and dimly lit laboratory. The stimulus images were presented on a 16-inch CRT monitor with a screen resolution of 1920 × 1080. Participants were required to complete facial expression identification tasks according to instructions presented on the monitor. Their electroencephalogram (EEG) data were collected during the experiment. For each trial, a focus point was presented for 1,000 ms. Subsequently, a facial image was presented, and the participant was required to determine whether the face was happy or angry by pressing a button (happy = 1; angry = 2) within 1,000 ms. If a button was pressed within 1,000 ms, then the picture disappeared, and a blank screen was presented until the next picture appeared. If no button was pressed, then the picture disappeared after 1,000 ms, and a blank screen was presented until the next picture appeared. The duration of the blank screens varied randomly from 900 ms to 1,700 ms. Figure 2 shows the experimental procedure. The experiment was divided into two blocks, each with 100 trials. The pictures within each block were balanced. Participants were given 2-3 min to rest between blocks.

**Figure 2.**
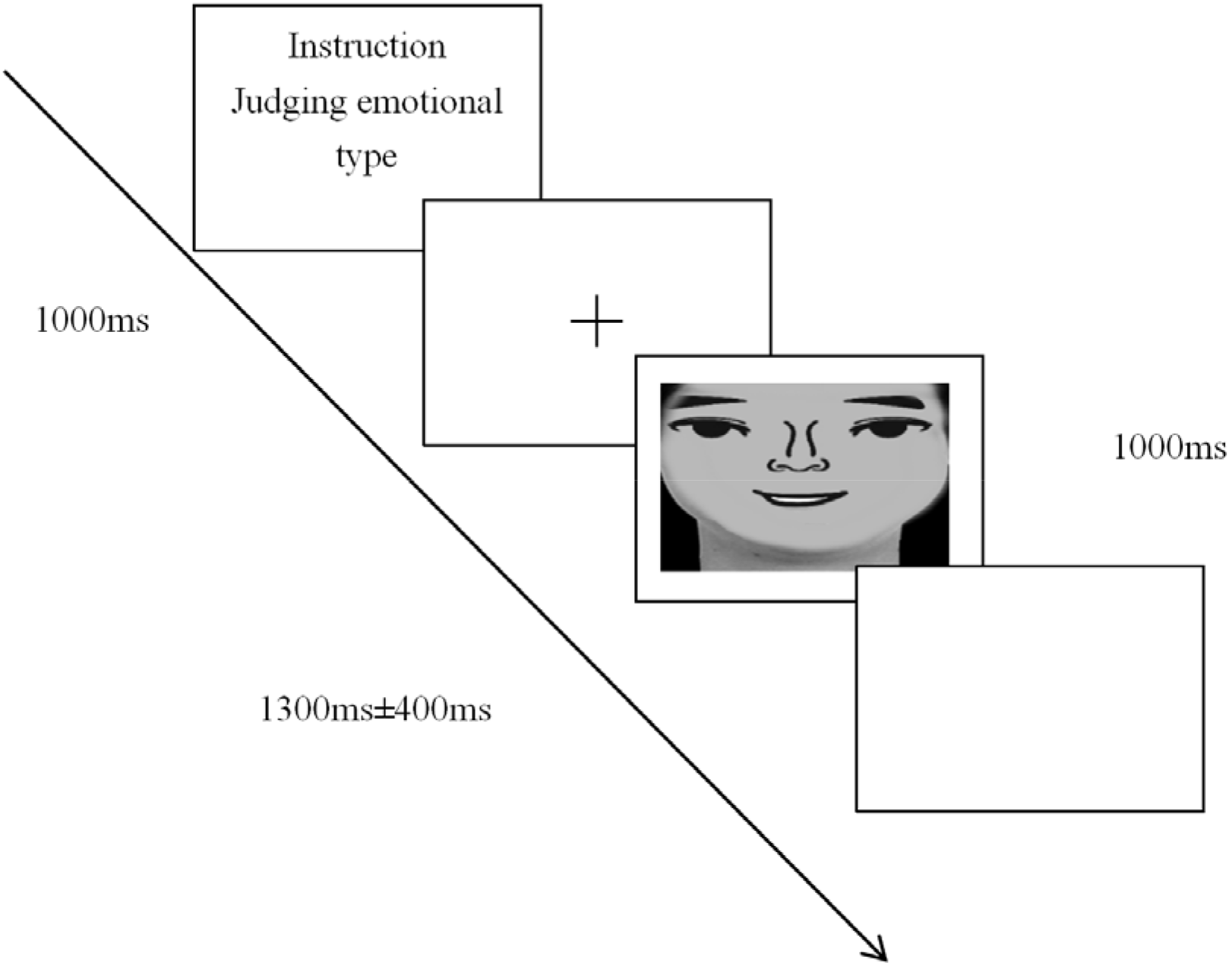
Task timing and example stimulus

### ERP measurement and data processing

We used a 64-lead EEG/ERP system (Electrical Geodesics, Inc., U.S.) for measurement and a potassium chloride solution as a conductive medium (skin impedance < 70 KΩ, A/D conversion = 24 bits, input impedance = 200 MΩ, sampling rate = 20 kHz, sampling range = ± 200 mV, amplifier noise = 1.4 μVpp, common mode rejection ratio = 120 dB, bandwidth = 0 ~ 4,000 Hz). Based on the overall mean chart, the early ERP components (N170 and VPP) generated by the stimuli showed clear peaks. A time window of 125-195 ms was used to measure the ERP peak and peak latency data collected at electrode locations in the left and right hemispheres (P7 and P8). For LPP, the average amplitude was calculated using the occipital lobe (O1, OZ, and O2), central zone (C3, CZ, and C4), and parietal lobe (P3, PZ, and P4) using a time window of 450-650 ms. Data processing, artifact elimination, and correction were performed.

### Statistical analyses

The behavioral and EEG data were analyzed using IBM SPSS 19.0 for Windows. N170 was analyzed using a 2 (face type: cartoon vs. real face) x 2 (emotional valance: happiness vs. anger) x 2 (gender: male vs. female) x 2 (hemisphere: left vs. right) repeated-measures analysis of variance (ANOVA). For VPP, a 2 (face type: cartoon vs. real face) x 2 (emotional valence: happiness vs. anger) x 2 (gender: male vs. female) repeated-measures ANOVA was used. For LPP, a 2 (face type: cartoon vs. real face) x 2 (emotion valence: happiness vs. anger) x 2 (gender: male vs. female) x 3 (brain region: occipital vs. central area vs. parietal) x 3 (lateralization: left vs. middle vs. right) repeated measures ANOVA was used.

## Result

### Behavioral Performance

**Table 2.**
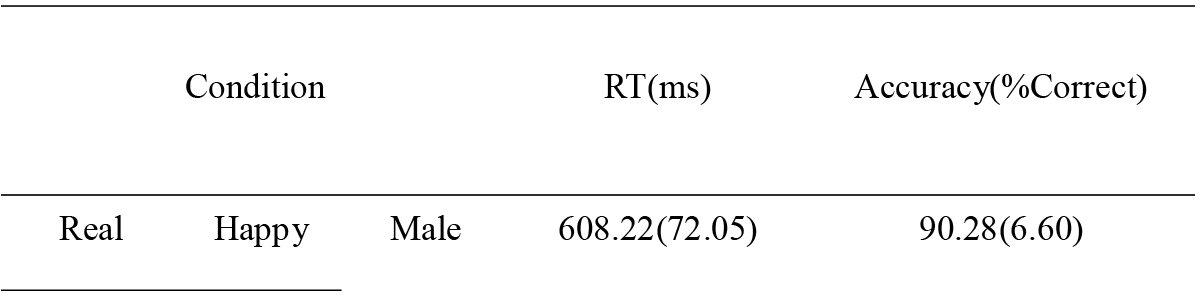
Means (and Standard Deviations) for Response Time (RT), Accuracy

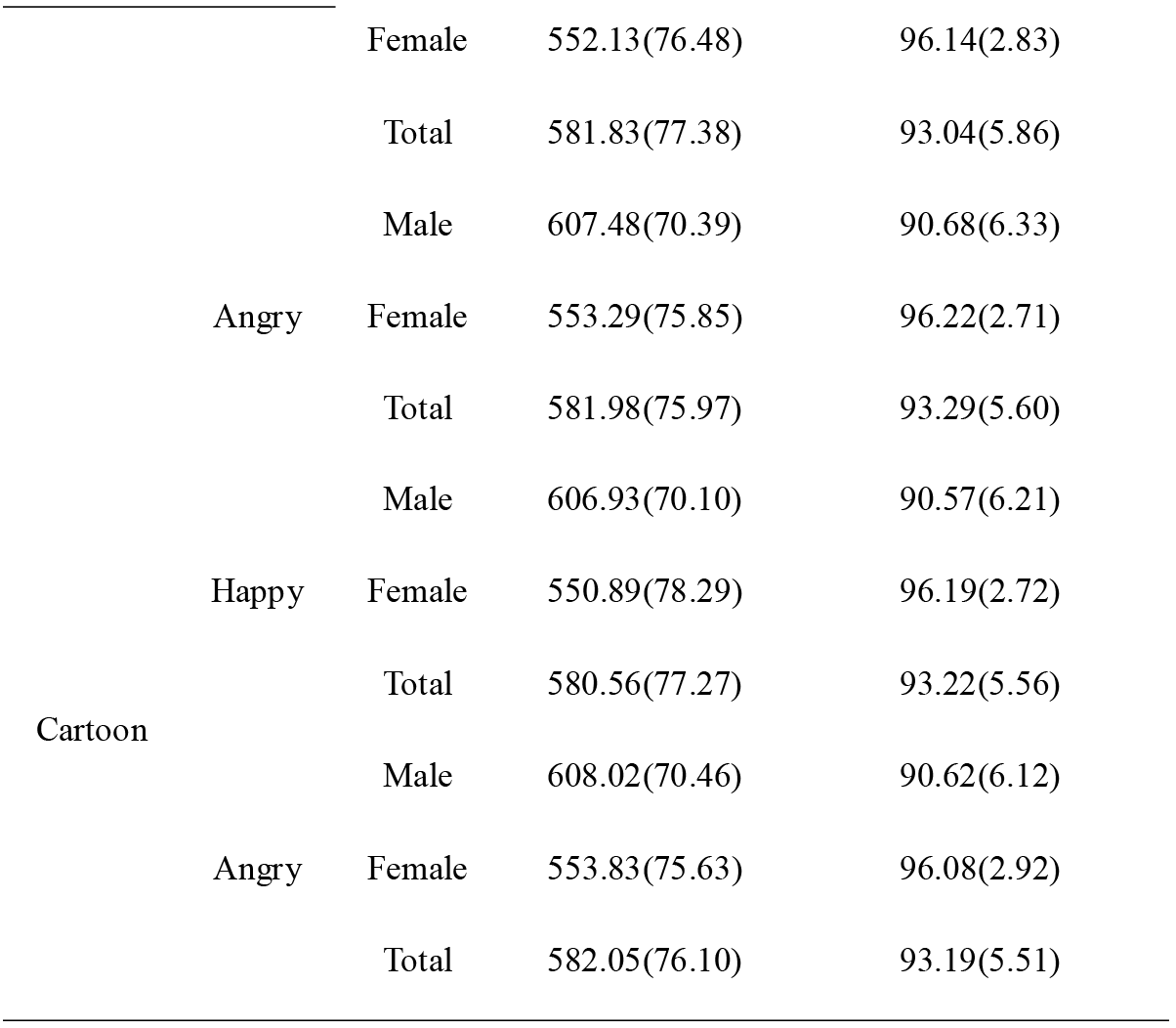

Mean response time (RT) and accuracy and standard deviations are shown in Table 2. Repeated-measures ANOVAs, with factors gender (male, female), face type (real, cartoon), emotion valence (happy, angry) as independent variables, with RT and accuracy as dependent variables, were conducted. For RT, a significant effect of emotion valence, F (1, 15) = 4.95, *p* < 0.05, η_p_^2^ = 0.25, revealed that RT was faster for happy face than angry face. The main effects of gender and face type did not reach significance, F (1, 15) = 2.40, *p* > 0.05, η_p_^2^ = 0.14, F (1, 15) = 2.17, *p* > 0.05, η_p_^2^ = 0.13, neither did all interactions (*ps* > 0.05).

For accuracy, a significant effect of gender, F (1, 15) = 5.38, *p* < 0.05, η_p_^2^ = 0.26, indicated that the female performed better than the male. Results also showed a significant interaction between gender and emotion valence, F (1, 15) = 5.50, *p* < 0.05, η_p_^2^ = 0.27. Follow-up simple effect analysis showed that for the male, performance was better in condition of angry face than happy face, F (1, 15) = 10.48, *p* < 0.01, η_p_^2^ = 0.41; for the female, accuracy did not differ for happy face versus angry face, F (1, 15) = 0.03, *p* > 0.05, η_p_^2^ = 0.002. The main effects of face type and emotion valence did not reach significance, F (1, 15) = 0.34, *p* > 0.05, η_p_^2^ = 0.02, F (1, 15) = 4.40, *p* > 0.05, η_p_^2^ = 0.23, neither did other interactions (*ps* > 0.05).

### N170

Mean amplitudes and latency of N170 and VPP and standard deviations are shown in Table 3. Repeated-measures ANOVAs, with factors gender (male, female), face type (real, cartoon), emotion valence (happy, angry), lateralization (P7 for left hemisphere, P8 for right hemisphere) as independent variables, with amplitudes and latency as dependent variables, were conducted. For amplitudes, a significant effect of face type, F (1, 13) = 34.77, *p* < 0.01, η_p_^2^ = 0.73, revealed that amplitudes were bigger for cartoon face than real face. N170 mean amplitudes from electrode P7 and P8 are shown in Figure 3. Results also showed a significant interaction between face type and lateralization, F (1, 13) = 6.43, *p* < 0.05, η_p_^2^ = 0.33. Differences between the amplitudes elicited by cartoon and real faces were bigger in right hemisphere (2.98±0.47μV) than left hemisphere (1.25±0.52μV). The main effects of gender, emotion valence and lateralization did not reach significance, F (1, 13) = 0.14, *p* > 0.05, η_p_^2^ = 0.01, F (1, 13) = 0.97, *p* > 0.05, η_p_^2^ = 0.07, F (1, 13) = 1.85, *p* > 0.05, η_p_^2^ = 0.12, neither did other interactions (*ps* > 0.05).

**Table 3.** Means (and Standard Deviations) for Amplitudes and Latency of N170 and VPP

| Condition | N170-Amplitude($\mu V$) | | N170-Latency(ms) | | VPP-Amplitude($\mu V$) | VPP-Latency(ms) |
| --- | --- | --- | --- | --- | --- | --- |
|  | P7 | P8 | P7 | P8 |  |  |

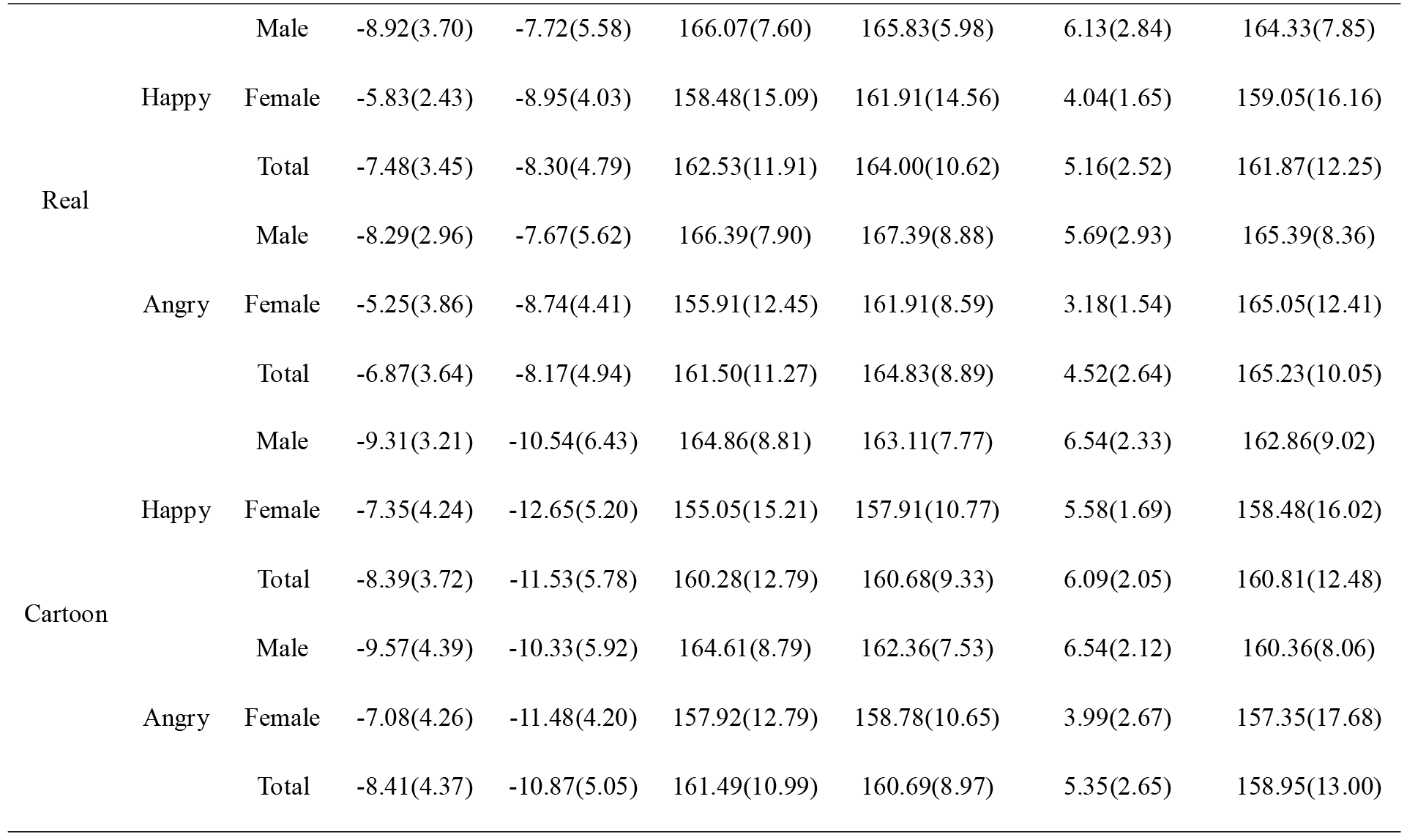

**Figure 3.**
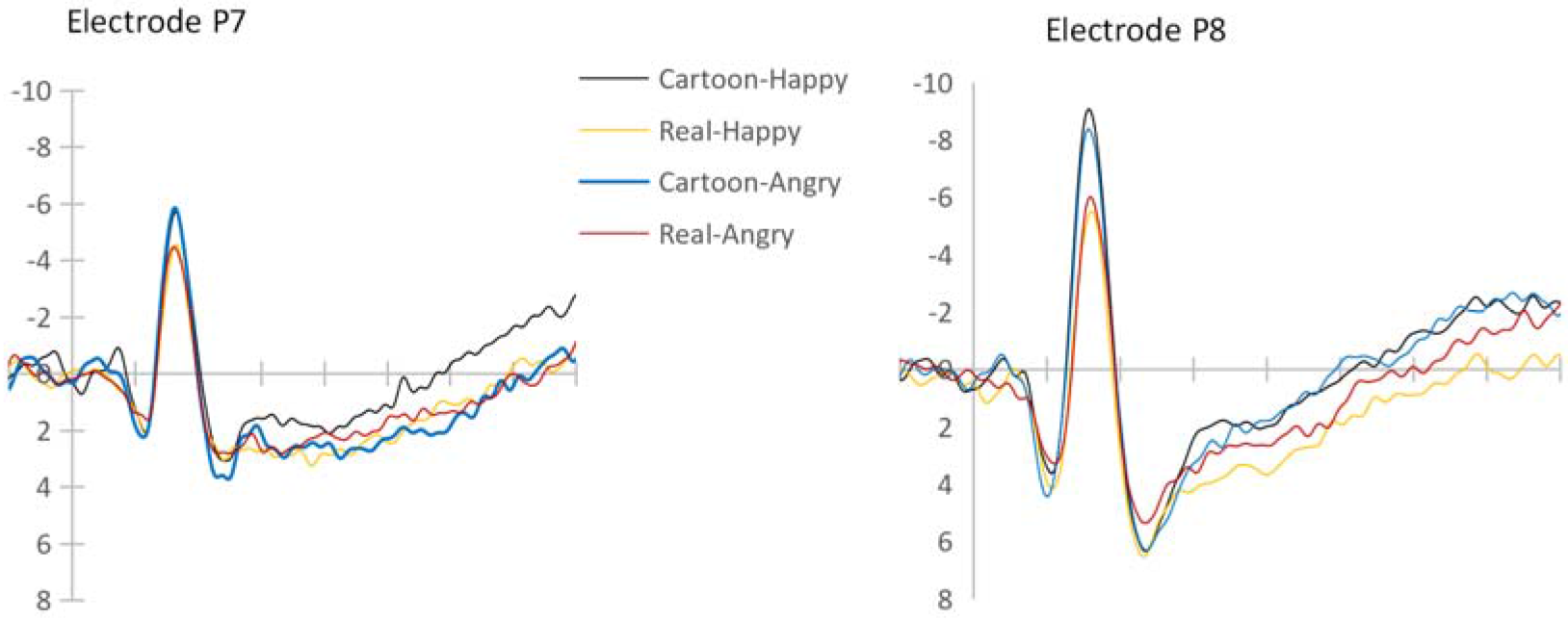
N170 mean amplitudes from electrode P7 and P8

For latency, a significant effect of face type, F (1, 13) = 15.16, *p* < 0.01, η_p_^2^ = 0.54, indicated that latency was longer for real face than cartoon face. Results also showed a significant interaction between face type and lateralization, F (1, 13) = 5.22, *p* < 0.05, η_p_^2^ = 0.29. Follow-up simple effect analysis showed that for right hemisphere, latency was longer in condition of real face than cartoon face, F(1, 13) = 26.69, *p* < 0.01, η_p_^2^ = 0.67; for left hemisphere, latency did not differ between real face and cartoon face, F (1, 13) = 1.35, *p* > 0.05, η_p_^2^ = 0.09. The main effects of gender, emotion valence and lateralization did not reach significance, F (1, 13) = 2.58, *p* > 0.05, η_p_^2^ = 0.17, F (1, 13) = 0.07, *p* > 0.05, η_p_^2^ = 0.01, F (1, 13) = 0.17, *p* > 0.05, η_p_^2^ = 0.01, neither did other interactions (*ps* > 0.05).

### VPP

Repeated-measures ANOVAs, with factors gender (male, female), face type (real, cartoon), emotion valence (happy, angry) as independent variables, with amplitudes and latency as dependent variables, were conducted. For amplitudes, a significant effect of face type, F (1, 13) = 4.94, *p* < 0.05, η_p_^2^ = 0.28, revealed that amplitudes were bigger for cartoon face than real face. The main effects of gender and emotion valence did not reach significance, F (1, 13) = 4.44, *p* > 0.05, η_p_^2^ = 0.26, F (1, 13) = 3.33, *p* > 0.05, η_p_^2^ = 0.20, neither did all interactions (*ps*>0.05).

For latency, results showed that interaction between face type and emotion valence was significant, F (1, 13) = 6.28, *p* < 0.05, η_p_^2^ = 0.33. Follow-up simple effect analysis found no significant effect. The main effects of gender, face type and emotion valence did not reach significance, F (1, 13) = 0.39, *p* > 0.05, η_p_^2^ = 0.03, F (1, 13) = 1.33, *p* > 0.05, η_p_^2^ = 0.09, F (1, 13) = 0.46, *p* > 0.05, η_p_^2^ = 0.03, neither did other interactions (*ps* > 0.05).

### LPP

Repeated-measures ANOVAs, with factors gender (male, female), face type (real, cartoon), emotion valence (happy, angry), braion region (top, central, occipital), lateralization (left, central, right) as independent variables, with amplitudes as dependent variables, were conducted. A significant effect of emotion valence, F (1, 13) = 4.38, *p* < 0.05, η_p_^2^ = 0.25, revealed that amplitudes were bigger for angry face (1.68±0.40μV) than happy face (1.02±0.52μV). Results showed a significant effect of face type, F (1, 13) = 6.47, *p* < 0.05, η_p_^2^ = 0.33, revealing that amplitudes were bigger for real face (2.02±0.49μV) than cartoon face (0.69±0.52μV). Another significant effect of lateralization, F (2, 12) = 10.38, *p* < 0.01, η_p_^2^ = 0.63, revealed that amplitudes were bigger for central (2.13±0.49μV) than left (0.72±0.51μV), and that amplitudes were bigger for right (1.21± 0.56μV) than left. A significant effect of region, F (2, 12) = 12.23, *p* < 0.01, η_p_^2^ = 0.67, revealed that amplitudes were bigger for central area (1.68±0.40μV) than occipital area (1.02±0.52μV), and that amplitudes were bigger for parietal area (1.82±0.46μV) than occipital area. Results also showed a significant interaction between region and lateralization, F (2, 12) = 5.20, *p* < 0.05, η_p_^2^ = 0.15. Follow-up simple effect analysis showed that for occipital area, amplitudes were bigger in condition of middle area than left and right hemispheres; for parietal area, amplitudes did not differ between hemispheres; for central area, amplitudes were bigger in condition of middle area and right hemisphere than left hemisphere. LPP mean amplitudes over central, occipital and parietal electrode are shown in Figure 4.

**Figure 4.**
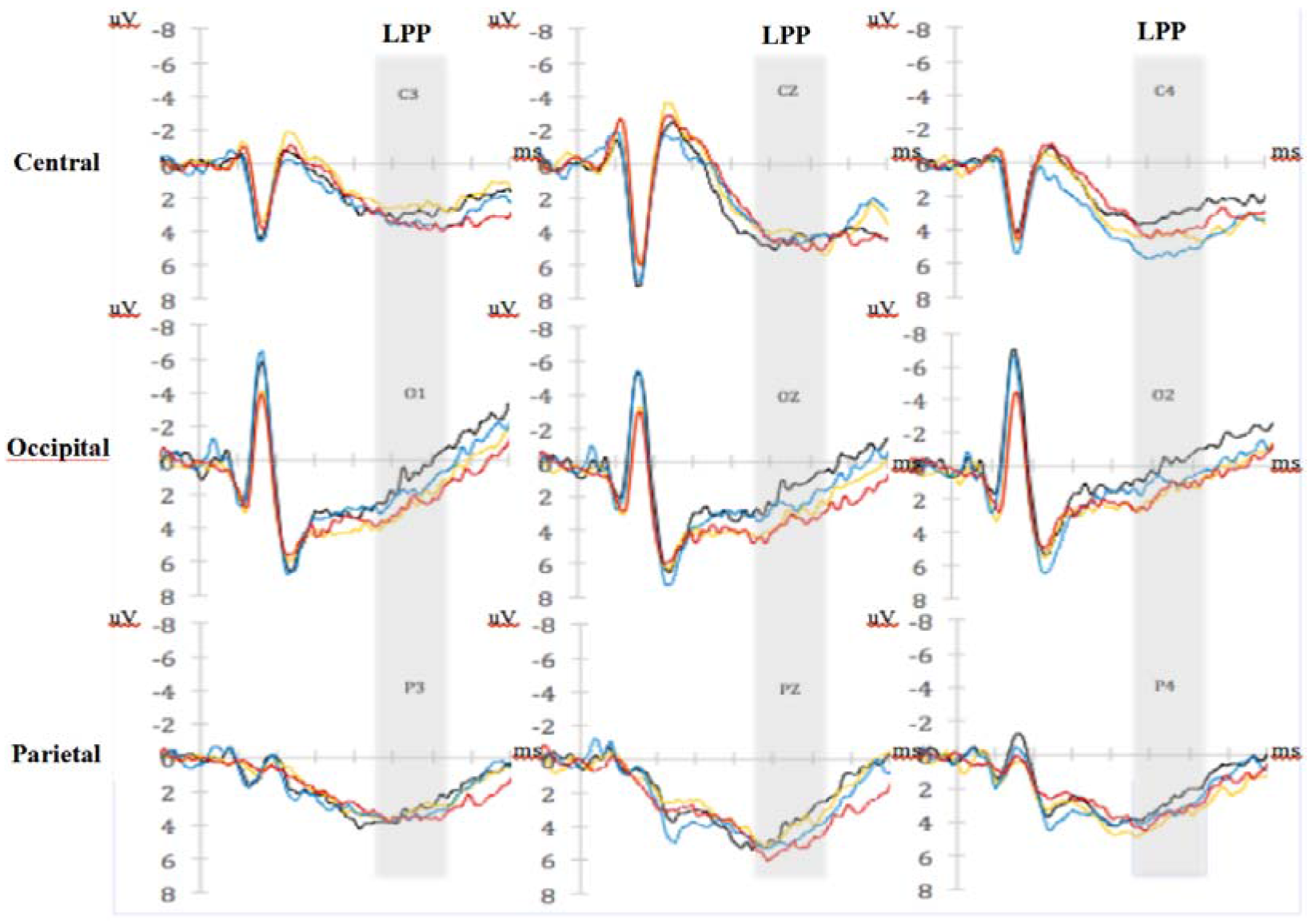
LPP mean amplitudes over central, occipital and parietal electrode

## Discussion

### Processing differences between cartoon and real faces

The differences in the processing of the two face types were primarily reflected by the amplitudes and latencies of N170, VPP, and LPP. Although no significant difference was found with regard to reaction time, cartoon faces resulted in shorter N170 latencies than did real faces. This finding is not consistent with previous studies. Wang et al. (2012) found that adults’ N170 latencies were shorter when viewing real faces than cartoon faces. These inconsistent results might have been caused by the differences in stimulus materials. In Wang’s study, real faces were collected from kindergarten children (mean age = ~6 years), and cartoon faces were obtained from screenshots of high-resolution DVD cartoons. In the present study, the real face stimuli were collected from adult facial expression databases (CFAPS and JAFFE), and the cartoon faces were converted from these real faces using MYOTee. The own-age bias (OAB, Wright & Stroud, 2002) might have caused these findings to differ from those of previous studies. The OAB states that people show better performance when recognizing faces from their own age group compared with other age groups. While both studies used adult participants, the age of viewed faces was closer to the participant age group in the present study.

A difference was also found in the N170 and VPP amplitudes during the early processing of real and cartoon faces. Our findings suggest a significant difference in brain processing intensity for the two face types during early processing. Cartoon faces were associated with significantly higher amplitudes than real faces. Previous studies (Wang et al., 2012; Sagiv & Bentin, 2001) have indicated that real faces induce larger N170 and VPP amplitudes than do cartoon faces. The current different result might be because of the different levels of abstraction of the cartoon faces in the different studies. Schindler et al. (2017) employed six face-stylization levels varying from abstract to realistic and investigated the difference in the processing of real and cartoon faces. The results revealed a U-shape relationship between N170 and face realism. That is, both the most abstract and most realistic faces caused stronger reactions compared with medium-stylized faces. The cartoon faces used in this study were converted from real faces using MYOTee. They are more simplified and abstract and, therefore, might have resulted in stronger N170 amplitudes. In addition, Proverbio, Riva, Martin and Zani (2010) found that infant faces elicited higher N170 amplitudes than did adult faces, most likely because of juvenile characteristics such as the larger proportion of the eyes. In the present study, the eyes of the cartoon faces were much larger than those of real people (including infants).

Real and cartoon faces also differed in LPP amplitude. The LPP amplitudes induced by real faces were significantly larger than those induced by cartoon faces. This finding is consistent with those of previous studies. When the neutral expressions of real faces and puppet faces were compared, no differences in N170 were observed. However, a stronger LPP was found with regard to real faces starting at 400 ms (Ma, Qian, Hu & Wang, 2017). This effect is because of the significance and uniqueness of the real face as well as the understanding of the portrayed individual (Wheatley, Weinberg, Looser, Moran & Hajcak, 2011) because computer-generated faces are usually more difficult to remember (Balas & Pacella, 2015; Crookes et al., 2015). Bruce and Young (1986) considered facial feature encoding and identify recognition as the second stage of face recognition. This stage includes the accurate processing of facial information such as age, gender, race, and facial expression. Adults invest more psychological resources to real faces during late face processing. Compared with simplified cartoon faces, real faces convey more personal information and social meaning. LPP is related to facial attractiveness (Ma et al., 2017; Marzi & Viggiano, 2010; Werheid, Schacht & Sommer, 2007). Therefore, the results of the present study might suggest that real faces are more attractive than simplified cartoon faces to adults.

Significant differences were found among various brain regions. LPP amplitudes in the central and parietal areas were greater than those in the occipital area, indicating that the central and parietal areas are the major brain regions for face processing, which is consistent with Bauer’s two-route model (1984). This model assumes that two routes exist for face recognition: the ventral route and the dorsal route. The dorsal route is primarily responsible for detecting the meaning of a face, beginning in the visual cortex, passing through the upper temporal sulcus, lower parietal lobe, and cingulate gyrus, eventually reaching the limbic system. In addition, the LPP amplitudes in the middle area and the right hemisphere were higher than those in the left hemisphere, suggesting a right hemisphere advantage. With regard to the N170 component, cartoon faces caused significantly smaller latencies than did real faces in the right hemisphere. However, no difference was observed in the left hemisphere. This finding is consistent with previous studies reporting a right hemisphere advantage with regard to face recognition intensity and speed (Yovel, Levy, Grabowecky & Paller, 2003). Furthermore, cartoon face processing showed obvious lateralization, primarily in the right hemisphere, whereas real face processing is bilateral (Wang et al., 2012).

### Differences between angry and happy faces in face processing

The ERP results revealed that LPP amplitudes distinguished emotion type during late processing. Angry faces induced larger positive LPP waves than did happy faces. This finding is consistent with Zhu and Liu (2014), who found that negative facial expressions were associated with significantly larger LPP amplitudes than were positive and neutral expressions. Previous studies (Holmes, Green & Vuilleumier, 2005; Whalen et al., 1998; Bradley, Mogg & Lee, 1997) reported bias in individual attention allocation when recognizing different facial expressions, especially those of negative emotions. Facial expression type did not influence N170. According to the face processing model of Bruce and Young (1986), facial expression processing is independent from facial structural processing. That is, facial emotion information should not influence N170. The research of Eimer et al. (2003) and Ashley, Vuilleumier snd Swick (2004) support this hypothesis.

### Differences between men and women with regard to facial expression processing

Women showed a higher facial expression recognition accuracy than did men. Extensive research (Hall, Hutton & Morgan, 2010; Brewster, Mullin, Dobrin & Steeves, 2011; Mcbain, Norton & Chen, 2009; Megreya, Bindemann & Havard, 2011) has demonstrated that women have a face recognition advantage over men. In addition, an interaction effect was found between facial expression type and gender. Men showed a higher accuracy for recognizing angry faces, whereas women showed no difference between the two expression types. Previous studies did not find this interaction. Overall, the recognition accuracy of happy faces was higher for both genders compared with angry faces. The high accuracy of angry face recognition among men might be because they are more physically aggressive than women (Glascock, 2008; Eagly & Steffen, 1986). High aggressiveness is positively correlated with serum testosterone concentration (Hu & Li, 2014). Therefore, men are likely more sensitive to social signals that convey aggressiveness.

### Advantages and limitations

Our selected stimuli (i.e., the cartoon and real facial expression pictures) represent the advantage of this study. One important advantage is that the cartoon faces were converted from real faces and therefore retain the same facial structure and hairstyle. Most of the existing research has used screenshots of cartoon characters or sketched faces and expression icons (Wang et al., 2012; Eastwood et al., 2001; Sagiv & Bentin, 2001); therefore, they cannot exclude nuanced information other than facial expressions compared with real faces. Another advantage is that all of the images used were of Asian adults, which prevented the introduction of cultural and age differences that might have been caused by use of western emotional faces. One limitation of this study is its limited sample size, which might have resulted in the large standard deviation in VPP latency. Although the interaction effect of face type x emotion valence was significant, the simple effects were not. Future research should increase the sample size to examine the interaction of cartoon and real faces with regard to facial expression type. Another limitation is that although ERP is advantageous for its temporal resolution, its special resolution is low. Future research should apply fMRI, which has a high spatial resolution. Finally, children are the primary audience for cartoons. Childhood is an important stage for developing emotional cognition. Based on the present study, future studies should compare adults and children with regard to the processing of cartoon and real facial expressions. This line of research might help draw a clearer picture of the developmental process associated with cartoon face processing.

### Conclusions

We used ERPs to measure the brain activity responses induced by the facial expressions of cartoon and real faces. According to the neurophysiological evidence in this study, face type has a strong but heterogeneous effect on the N170, VPP, and LPP components. During the early processing stage, adults process cartoon faces faster than real faces. However, adults allocate more attentional resources for real face processing during late processing stage. Facial expression type influenced late-stage component LPP, showing attentional bias for negative emotions; however, early-state N170 was not influenced. Future research should use larger sample sizes to examine the interaction between face type (real vs. cartoon) and facial expression.

## References

Ashley, V., Vuilleumier, P., & Swick, D. (2004). Time course and specificity of event-related potentials to emotional expressions. Neuroreport, 15(1), 211–216.

Balas, B., & Pacella, J. (2015). Artificial faces are harder to remember. Computers in Human Behavior, 52, 331–337.

Batty, M., & Taylor, M. J. (2003). Early processing of the six basic facial emotional expressions. Brain Research Cognitive Brain Research, 17(3), 613.

Bauer, R. M. (1984). Autonomic recognition of names and faces in prosopagnosia: a neuropsychological application of the guilty knowledge test. Neuropsychologia, 22(4), 457–469.

Bentin, S., Allison, T., Puce, A., Perez, E., & Mccarthy, G. (1996). Electrophysiological studies of face perception in humans. Journal of Cognitive Neuroscience, 8(6), 551–565.

Bernat, E., Bunce, S., & Shevrin, H. (2001). Event-related brain potentials differentiate positive and negative mood adjectives during both supraliminal and subliminal visual processing. International Journal of Psychophysiology Official Journal of the International Organization of Psychophysiology, 42(1), 11–34.

Bradley, B. P., Mogg, K., & Lee, S. C. (1997). Attentional biases for negative information in induced and naturally occurring dysphoria. Behaviour Research & Therapy, 35(10), 911–27.

Bruce, V., & Young, A. (1986). Understanding face recognition. British Journal of Psychology, 77(3), 305.

Bublatzky, F., Gerdes, A. B., White, A. J., Riemer, M., & Alpers, G. W. (2014). Social and emotional relevance in face processing: happy faces of future interaction partners enhance the late positive potential. Frontiers in Human Neuroscience, 8, 1–10.

Calvo, M. G., & Lundqvist, D. (2008). Facial expressions of emotion (kdef): identification under different display-duration conditions. Behav Res Methods, 40(1), 109–115.

Chen, H., Russell, R., Nakayama, K., & Livingstone, M. (2010). Crossing the ‘uncanny valley’: adaptation to cartoon faces can influence perception of human faces. Perception, 39(3), 378–386.

Codispoti, M., Ferrari, V., & Bradley, M. M. (2006). Repetitive picture processing: autonomic and cortical correlates. Brain Research, 1068(1), 213–220.

Crookes, K., Ewing, L., Gildenhuys, J. D., Kloth, N., Hayward, W. G., & Oxner, M., et al. (2015). How well do computer-generated faces tap face expertise?. Plos One, 10(11), e0141353.

Eagly, A. H., & Steffen, V. J. (1986). Gender and aggressive behavior: a meta-analytic review of the social psychological literature. Psychological Bulletin, 100(3), 309–330.

Eastwood, J. D., Smilek, D., & Merikle, P. M. (2001). Differential attentional guidance by unattended faces expressing positive and negative emotion. Perception & Psychophysics, 63(6), 1004–1013.

Ekman, P., & Friesen, W. V. (1978). Facial action coding system (facs): a technique for the measurement of facial actions. Rivista Di Psichiatria,47(2), 126–38.

Ekman, P., & Friesen, W. V. (1971). Constants across cultures in the face and emotion. Journal of Personality & Social Psychology, 17(2), 124–129.

Eimer, M., Holmes, A., & Mcglone, F. P. (2003). The role of spatial attention in the processing of facial expression: an erp study of rapid brain responses to six basic emotions. Cognitive Affective & Behavioral Neuroscience, 3(2), 97–110.

Erickson, K., & Schulkin, J. (2003). Facial expressions of emotion: a cognitive neuroscience perspective. Brain & Cognition, 52(1), 52–60.

Flaisch, T., Häcker, F., Renner, B., & Schupp, H. T. (2011). Emotion and the processing of symbolic gestures: an event-related brain potential study. Social Cognitive & Affective Neuroscience, 6(1), 109.

Galli, G., Feurra, M., & Viggiano, M. P. (2006). “did you see him in the newspaper?” electrophysiological correlates of context and valence in face processing. Brain Research, 1119(1), 190–202.

Hajcak, G., Moser, J. S., & Simons, R. F. (2006). Attending to affect: appraisal strategies modulate the electrocortical response to arousing pictures. Emotion, 6(3), 517–522.

Hansen, C. H., & Hansen, R. D. (1988). Finding the face in the crowd: an anger superiority effect. Journal of Personality & Social Psychology,54(6), 917–24.

Han, S., Gao, X., Humphreys, G. W., & Ge, J. (2008). Neural processing of threat cues in social environments. Human Brain Mapping, 29(8), 945.

Hietanen, J. K., & Astikainen, P. (2013). N170 response to facial expressions is modulated by the affective congruency between the emotional expression and preceding affective picture. Biological Psychology, 92(2), 114–24.

Hinojosa, J. A., Mercado, F., & Carretié, L. (2015). N170 sensitivity to facial expression: a meta-analysis. Neuroscience & Biobehavioral Reviews, 55, 498–509.

Hoffmann, H., Kessler, H., Eppel, T., Rukavina, S., & Traue, H. C. (2010). Expression intensity, gender and facial emotion recognition: women recognize only subtle facial emotions better than men. Acta Psychologica, 135(3), 278–283.

Holmes, A., Green, S., & Vuilleumier, P. (2005). The involvement of distinct visual channels in rapid attention towards fearful facial expressions. Cognition & Emotion, 19(6), 899–922.

Hoptman, M. J., & Levy, J. (1988). Perceptual asymmetries in left- and right-handers for cartoon and real faces. Brain & Cognition, 8(2), 178–188.

Hu, B. P., & Li, S. J. (2014). Studies on Serotonin, Testosterone and Total Cholesterol in the Blood of Highly Aggressive People. Journal of Physiology Studies, 02, 13–18.

Jack Glascock Ph.D. (2008). Direct and indirect aggression on prime-time network television. Journal of Broadcasting & Electronic Media,52(2), 268–281.

Jessica K. Hall, Sam B. Hutton, & Michael J. Morgan. (2010). Sex differences in scanning faces: does attention to the eyes explain female superiority in facial expression recognition?. Cognition & Emotion, 24(4), 629–637.

Joyce, C., & Rossion, B. (2005). The face-sensitive n170 and vpp components manifest the same brain processes: the effect of reference electrode site. Clinical Neurophysiology, 116(11), 2613–2631.

Keil, A., Bradley, M. M., Hauk, O., Rockstroh, B., Elbert, T., & Lang, P. J. (2002). Large-scale neural correlates of affective picture processing. Psychophysiology, 39(5), 641–649.

Kendall, L. N., Raffaelli, Q., Kingstone, A., & Todd, R. M. (2016). Iconic faces are not real faces: enhanced emotion detection and altered neural processing as faces become more iconic. Cognitive Research Principles & Implications, 1(1), 19.

Krolak-Salmon, P., Fischer, C., Vighetto, A., & Mauguière, F. (2001). Processing of facial emotional expression: spatio-temporal data as assessed by scalp event-related potentials. European Journal of Neuroscience, 13(5), 987.

Ma, Q., Qian, D., Hu, L., & Wang, L. (2017). Hello handsome! male’s facial attractiveness gives rise to female’s fairness bias in ultimatum game scenarios-an erp study. Plos One, 12(7), e0180459.

Marzi, T., & Viggiano, M. P. (2010). When memory meets beauty: insights from event-related potentials. Biological Psychology, 84(2), 192–205.

Mcbain, R., Norton, D., & Chen, Y. (2009). Females excel at basic face perception. Acta Psychologica, 130(2), 168.

Megreya, A. M., Bindemann, M., & Havard, C. (2011). Sex differences in unfamiliar face identification: evidence from matching tasks. Acta Psychologica, 137(1), 83–89.

Mullin, C. R. (2011). Sex differences in face processing are mediated by handedness and sexual orientation. Laterality, 16(2), 188–200.

Nelson, C. A., Morse, P. A., & Leavitt, L. A. (1979). Recognition of facial expressions by seven-month-old infants. Child Dev, 50(4), 1239–1242.

Proverbio, A. M., Riva, F., Martin, E., & Zani, A. (2010). Face coding is bilateral in the female brain. Plos One, 5(6), e11242.

Recio, G., Sommer, W., & Schacht, A. (2011). Electrophysiological correlates of perceiving and evaluating static and dynamic facial emotional expressions. Brain Research, 1376(3), 66.

Rellecke, J., Sommer, W., & Schacht, A. (2012). Does processing of emotional facial expressions depend on intention? time-resolved evidence from event-related brain potentials. Biological Psychology,90(1), 23–32.

Rossion, B., & Jacques, C. (2008). Does physical interstimulus variance account for early electrophysiological face sensitive responses in the human brain? ten lessons on the n170. Neuroimage, 39(4), 1959–1979.

Sagiv, N., & Bentin, S. (2001). Structural Encoding of Human and Schematic Faces: Holistic and Part-Based Processes. MIT Press.

Schindler, S., & Kissler, J. (2016). People matter: perceived sender identity modulates cerebral processing of socio-emotional language feedback. Neuroimage, 134(2), 160–169.

Schindler, S., Zell, E., Botsch, M., & Kissler, J. (2017). Differential effects of face-realism and emotion on event-related brain potentials and their implications for the uncanny valley theory. Scientific Reports, 7, 45003.

Schupp, H. T., Flaisch, T., Stockburger, J., & Junghöfer, M. (2006). Emotion and attention: event-related brain potential studies. Progress in Brain Research, 156(156), 31–51.

Schupp, H. T., Junghöfer, M., Weike, A. I., & Hamm, A. O. (2004). The selective processing of briefly presented affective pictures: an erp analysis. Psychophysiology, 41(3), 441–449.

Steppacher, I., Schindler, S., & Kissler, J. (2015). Higher, faster, worse? an event-related potentials study of affective picture processing in migraine. Cephalalgia An International Journal of Headache, 36(3), 249.

Thompson, D. F., & Meltzer, L. (1964). Communication of emotional intent by facial expression. Journal of Abnormal Psychology, 68(2), 129.

Wang, L.,Wang, J. M., Wang, J. L., & Lu, Y. J. (2012). A Comparative Study of Event-related Potential in Cartoon Faces and Real Faces Recognition. Psychological Research, 5(5), 19–28.

Wang, Y., & Luo, Y. J. (2005). Standardization and Evaluation of College Students’ Facial Expression Materials. Chinese Journal of Clinical Psychology, 13(4), 396–398.

Wei, J. H., & Luo, Y. J. (2002). Event-related Brain Potentials: The Cognitive ERP Textbook. Beijing: The Economic Daily Press, 2002, 5:8–32.

Werheid, K., Schacht, A., & Sommer, W. (2007). Facial attractiveness modulates early and late event-related brain potentials. Biological Psychology, 76(2), 100–108.

Whalen, P. J., Rauch, S. L., Etcoff, N. L., Mcinerney, S. C., Lee, M. B., & Jenike, M. A. (1998). Masked presentations of emotional facial expressions modulate amygdala activity without explicit knowledge. Journal of Neuroscience, 18(1), 411–418.

Wheatley, T., Weinberg, A., Looser, C., Moran, T., & Hajcak, G. (2011). Mind perception: real but not artificial faces sustain neural activity beyond the n170/vpp. Plos One, 6(3), e17960.

Wieser, M. J., Pauli, P., Reicherts, P., & Mühlberger, A. (2010). Don’t look at me in anger! enhanced processing of angry faces in anticipation of public speaking. Psychophysiology, 47(2), 271–280.

Wildgruber, D., Pihan, H., Ackermann, H., Erb, M., & Grodd, W. (2002). Dynamic brain activation during processing of emotional intonation: influence of acoustic parameters, emotional valence, and sex. Neuroimage, 15(4), 856–869.

Wright, D. B., & Stroud, J. N. (2002). Age differences in lineup identification accuracy: people are better with their own age. Law & Human Behavior, 26(6), 641–654.

Yovel, G., Levy, J., Grabowecky, M., & Paller, K. A. (2003). Neural correlates of the left-visual-field superiority in face perception appear at multiple stages of face processing. Cognitive Neuroscience Journal of,15(3), 462–474.

Zhu, Y. Y., & Liu, Z. X. (2014). An ERP Study of Dynamic Facial Expression Recognition under Different Attention Conditions. Chinese Journal of Applied Psychology, 20(4), 375–384.

